# Comparative sensitivity of the test with tuberculosis recombinant allergen, containing ESAT6-CFP10 protein, and Mantoux test with 2 TU PPD-L in newly diagnosed tuberculosis children and adolescents in Moscow

**DOI:** 10.1101/343665

**Authors:** Liudmila Slogotskaya, Elena Bogorodskaya, Diana Ivanova, Tatiana Sevostyanova

**Affiliations:** Clinical Research, Scientific and Clinical Antituberculosis Center of Moscow Government Departament of Clinical Research, Scientific and Clinical Antituberculosis Center of Moscow Government Health Department, Moscow; Russian Federation

## Abstract

**Background.** A group of Russian scientists has developed Diaskintest, which comprises *Mycobacterium tuberculosis*-specific recombinant proteins CFP10-ESAT6, for skin testing (0.2 µg/0.1 ml).

**Study purpose:** to evaluate the comparative sensitivity of TST with 2 TU PPD-L and a skin test with tuberculous recombinant allergen (Diaskintest) containing the ESAT6-CFP10 protein in children and adolescents with newly diagnosed respiratory tuberculosis during mass screening in the primary medical service in Moscow, 2013–2016.

**Materials and methods.** The trial was a comprehensive retrospective group study of children and adolescents diagnosed in Moscow with respiratory tuberculosis in 2013–2016, aged 0 to 17 years inclusive. From 441 patients selected for analysis 408 patients had both tests (TST with 2 TU PPD-L and Diaskintest) performed, in 193 patients both tests were given simultaneously, of them 162 patients were BCG-vaccinated.

**Results.** Comparative results of both tests in 408 patients with tuberculosis: at cut-off ≥ 5 mm, both tests has similar sensitivity: Diaskintest 98.3 % (95 % CI 97.0–99.6 %), TST 98.0 % (95 % CI 96.7–99.4 %), at cut-off ≥10 mm, the sensitivity decreases for both tests: Diaskintest 90.0 % (95 % CI 87.0–93.0 %), TST 88.7 % (95 % CI 85.6–91.9 %), but at cut-off ≥ 15 mm, the decrease in sensitivity is statistically significant: for Diaskintest 61.5 % (95 % CI 56.7–66.3 %), and for TST 46.3 % (95 % CI 41.4–51.3 %), p <0.0001.

The results of simultaneous setting of tests on different hands in 193 people (including 162 BCG-vaccinated), do not differ from the results for 408 people.

The correlation between the results of Diaskintest and TST was significant in all groups.

**Conclusion.** In children and adolescents with respiratory tuberculosis, Diaskintest of 0.2 µg/ml and the Mantoux test with 2 TU PPD-L have high sensitivity (98%) at a cut-off of 5 mm; however, at cut-off ≥ 15 mm sensitivity is significantly reduced, and the decrease is more pronounced in the Mantoux test. The advantage of Diaskintest is that, unlike the Mantoux test, it has high specificity under the conditions of mass BCG vaccination. The test is cost-effective, simple to carry out, and can be used in mass screening.

## Introduction

The diagnosis and treatment of latent tuberculosis infection (LTBI) is one of the strategies recommended by the World Health Organization (WHO) to combat tuberculosis (TB) worldwide and is part of the WHO strategy for tuberculosis control [1]. With regard to acceptable methods for diagnosing LTBI, WHO documents recommend that either a tuberculin test (TST) or an IGRA test be used to detect LTBI in high-income and middle-income countries and a TB incidence of <100 per 100,000 population. TST is preferred in low- and middle-income countries with an incidence of ≥100 per 100,000 population [2,3]. Due to the low specificity of the Mantoux test, the high frequency of false positives due to cross-sensitization with the BCG vaccine strain *(Mycobacterium bovis BCG)* – difficulties arise in interpreting it. [4,5].

An important stage in improving the methods of diagnosis of tuberculosis was the possibility of studying and deciphering the genome of tuberculosis mycobacteria, which allowed to identify the differences between *M. bovis BCG* vaccine strain and *Mycobacterium tuberculosis virulent strains.* In particular, in *M. tuberculosis* genome, a region of difference *(RDI)* was discovered, which contains genes that code for the secretion of CFP10 and ESAT6 proteins. These proteins are expressed on the surface of the mycobacterial cell upon its multiplication and determine the virulent properties of the mycobacterium. Such a site is absent in *M. bovis BCG* genome and most nontuberculous mycobacteria, which allows differentiating postvaccinal and infectious allergies [6,7,8]. The discovery of antigens specific for *M. tuberculosis* led to the development of *in vitro* tests based on the production of gamma-interferon (IFN-γ) in response to stimulation with these antigens (IGRA-Interferon-Gamma Release Assays) [8,9]. The tests showed almost 100 % specificity, but a lower sensitivity – about 80 % [10,11,12].

The results of IGRA tests in the children’s population are mixed, because the confirmation or exclusion of the local form of tuberculosis in children is a difficult task [13,14]. According to B. Kampmann and co-authors [15], the sensitivity indicators for active tuberculosis in children were from 58 % to 80 %. In their opinion, the tests do not identify the disease in a significant proportion of children. But negative test results of tuberculin and IGRA allow to exclude, with 95 % confidence, the presence of tuberculosis infection [16,17]. Interest in the diagnosis of tuberculosis in children is gaining momentum, in particular, study of new biomarkers that allow detecting tuberculosis infection [18].

IGRA tests are most useful in populations immunized with the BCG vaccine, since this is where their specificity comes in handy [19].

There is no international agreement on cut-off values for the definition of a positive tuberculin reaction [20]. The choice among commonly used cut-off values (e.g. a diameter of induration of ≥ 5mm, ≥ 10mm or ≥ 15mm) depends on an individual’s risk factor profile for TB. Usually, a lower cut-off value of ≥ 5mm is used for individuals at higher risk of TB and a higher cut-off value of ≥ 10mm is applied for individuals at lower risk of TB [21,22,23].

The above-mentioned limitations are compounded by issues related to the interpretation of test results, which may independently influence false-positive and false-negative rates of the TST (e.g. different cut-off values, PPD dose) [20,21,24].

In general, both pooled sensitivity and specificity values of the IGRAs and the TST were similarly high in people who were not vaccinated with BCG (> 90%); however, the pooled specificity of the TST in BCG-vaccinated populations was much lower than that of IGRAs (about 56% vs. 96%) [25,26, 27].

In contrast, prospective longitudinal studies showed that neither the IGRAs nor the TST had a high prognostic value in predicting the risk of progression to active TB [27, 28].

Sollai S. et al., [29] in their meta-analysis conducted a comparative analysis of the sensitivity of IGRA tests and Mantoux skin test in high and low income countries for the first time. In *high-income* countries, the QFT-G-IT sensitivity was 0.79 (95 % CI 0.75–0.82). In *low-income* countries, the sensitivity of QFT-G-IT was significantly lower: 0.57 (95 % CI 0.52–0.61. The higher specificity of IGRA versus TST was observed in high-income countries (97–98 % vs. 92 %), but not in low-income countries (85–93 % vs. 90 %).

Five meta-analyzes published earlier evaluated the sensitivity and specificity of IGRA in children, but the data on the overall assessment vary considerably. In them, similarly to the data presented in adults, IGRA was reported to be more specific in comparison with TST. However, the IGRA sensitivity was 62 % to 89 % for T-SPOT.TB and 66 % to 83 % for QFT-G-IT [26, 30-33].

Thus, the currently available information is insufficient to assess the sensitivity, specificity and reproducibility of IGRA tests in children.

IGRA tests, having high specificity, still have a number of serious disadvantages: high material costs, the need for an equipped laboratory, intravenous manipulation and the precautionary requirements for maintaining the viability of lymphocytes producing INF-γ, which does not allow this method to be used for mass diagnostics.

The solution of the problem for Russia was the introduction of a skin test with CFP10 and ESAT6 proteins. In Russia, the medicinal product Diaskintest^®^ (manufactured by Generium according to the GMP standard) was developed, which is a complex of CFP10/ESAT6 recombinant proteins, produced by *Escherichia coli, BL21 (DE3) / pCFP-ESAT*, intended for setting an intradermal test in a dose of 0.1 μg/0.1 ml according to the Mantoux test technique [34]. The test has been widely used in Russia since 2009 according to the order of the Ministry of Health of Russia.

Similar to the Russian test, the Danish skin test with C-Tb was developed, which is produced by *Statens Serum Institut* in accordance with the GMP standard (Copenhagen, Denmark) – the difference is that it is a mixture of two recombinant proteins ESAT-6 and CFP10 in the ratio of 1:1. Both proteins are produced by *Lactococcus lactis*, the dose of the drug for intradermal application is 0.1 μg/0.1 ml. The test underwent 3 phases of clinical trials [35-38].

In general, compared to TST based on PPD, IGRA tests and new skin tests Russian Diaskintest and Danish C-tb test may offer additional benefits, especially improved specificity [39]. Of course, the availability of new tools is a critical factor for expanding their application, these are cost-effectiveness and feasibility in conditions of a lack of resources [40].

Recent international reviews of WHO strategy END TB specify the priority tasks: development of *biomarkers* for the detection and diagnosis of tuberculosis in children and people from high-risk groups, including systematic *screening* (active search for TB cases); detection of *latent tuberculosis infection* – to reduce the pool of individuals with LTBI; evaluation of treatment effectiveness. A biomarker shall have low cost – for TB diagnostics at the primary healthcare level [18]. Since there is no diagnostic standard for LTBI, active tuberculosis is used as a comparable substitute, although tuberculosis is known to be immunosuppressive.

### Study purpose

study of the comparative sensitivity of Mantoux test with 2 TU PPD-L and a skin test with tuberculous recombinant allergen (Diaskintest) containing the ESAT6– CFP10 protein in newly diagnosed tuberculosis in children and adolescents registered in 2013– 2016 in Moscow in a mass application of tests in the primary medical service.

## Materials and methods

### Study designs

The design of the study – a full-design retrospective group study of children and adolescents diagnosed in Moscow with respiratory tuberculosis in 2013–2016.

In Russia, all children are to be vaccinated with BCG after birth and revaccinated at the age of 7 provided that they have a negative Mantoux test with 2 TU PPD-L. Every year in Moscow all children and adolescents are screened with the Mantoux test with 2TU PPD-L (Saint Petersburg RDE of Vaccines and Serums FMBA of Russia) in order to assess the presence of post-vaccination allergy and identify a changed sensitivity to tuberculin (a positive reaction after complete test extinction, an increase in the positive reaction of 6 mm or more). Every year in Moscow, all children and adolescents are subject to a scheduled Mantoux test with 2 TU PPD-L (TST) to detect a change in response to a tuberculin test (the appearance of a positive reaction, an increase in the positive response by 6 mm or more). A cut-off≥ 5 mm is considered for TST in Russia. Children and adolescents with a positive TST should be examined by Diaskintest (CJSC GENERIUM, Russia). Tuberculosis recombinant allergen (Diaskintest) is administered at a dose of 0.2 μg in 0.1 ml in Mantoux technique. When evaluating the response to the introduction of the tuberculosis recombinant allergen, the test was regarded as negative in the absence of infiltration or hyperemia, doubtful in the presence of only hyperemia of any size, positive in the presence of an infiltrate of any size.

Sometimes, at the request of parents, children are given two tests simultaneously on different hands.

Persons with a positive reaction to Diaskintest are subject to a thorough examination to identify local forms of tuberculosis (pulmonary and extrapulmonary). This is a computed tomography of chest organs, ultrasound of other organs, and in the presence of changes in the lungs – a sputum examination on MBT culture and molecular genetic methods. In the presence of local changes in the chest, including the intrathoracic lymph nodes and the pleura, children are subject to a commission assessment of changes in order to establish a diagnosis of tuberculosis or the presence of post-tuberculosis changes. The commission consists of the most qualified phthisiatricians and radiologists. If a diagnosis of tuberculosis is established, the children are hospitalized in a 24–hour in-patient facility and given anti-tuberculosis therapy, depending on the severity of the process, the availability of drug resistance, but at least 6 months. All antituberculosis activities are free, regardless of whether they are permanent Moscow residents or migrants. Children with positive reactions without local changes are subject to preventive therapy for at least 3 months on an outpatient basis or in a sanatorium and are observed during the year.

There was no formal sample size. The study was conducted as a full-design study.

### Main approaches to the statistical analyses

#### Descriptive statistics

The results contains the descriptive statistics of the subjects included to this study. For quantitative variables (age, the numeric results of the diagnostic tests, etc.) the following statistics presented: number of valid values (N); minimum (Min); maximum (Max); arithmetic mean (M); standard deviation (SD); 95% confidence interval (CI) for the mean; median (Me); interquartile range (IQR). Categorical data (such as gender, a group of the subject, etc.) presented with frequency in a format n/N and percentage (%).

#### Analyses of the test results

Data on sensitivity of the immunological tests presented as frequencies in a format n/N, where n is the number of subjects with true positive test results and N is number of subjects included to the analysis, as well as respective percentage (%).

McNemar test was applied to compare the results on sensitivity and concordance/disconcordance of the two tests and to estimate the odds ratios (OR) for a true positive result under different cut-off levels applied. To reveal chance-corrected agreement between the two tests, the kappa coefficients were calculated. The data presented with all the count a-b-c-d for two-by-two tables under different combinations of the test results.

Quantitative results of the two tests (size of indurations in mm) were analyzed with Pearson correlation and paired t-test. The point estimate for the mean differences in induration sizes along with 95% CI are shown.

#### Ethical approvals

The trial was approved by the Ethics Committee of Scientific and Clinical Antituberculosis Center of Moscow Government Health Department, Moscow (number 3, 17.11.2016) conducted in accordance with the principles of Good Clinical Practice and the World Medical Association (WMA) Declaration of Helsinki adopted by the 18th WMA General Assembly, Helsinki, Finland, 1964 and subsequent amendments.

All parents give written informed consent for carrying out skin tests and any manner of examination and treatment. The patients’ parents gave their consent to the processing of the study data provided that no personal information is published. Access to medical documents for the conduct of this study was granted on November 17, 2016.

#### Software

All the statistical analyses were performed using Stata ver. 14 (StataCorp LP, www.stata.com. Stata Statistical Software: Release 14. College Station T:SL, 2).

## Results and discussion

### Results

#### Analyses of baseline characteristics of the subjects

The following age category of subjects was included in the study: children and adolescents aged 0 to 17 years inclusive – total 441. Out of 441 patients, the results of the Diaskintest are known in 421 people, TST- in 414 people (the reasons of no TST results were: parents refusal to perform TST having only Diaskintest done; no data for patients who are not residents, although they were in the territory of Moscow, entered the official statistics of morbidity, received treatment in federal healthcare institutions and left for home, without submitting documents to the Moscow TB Service). The mean age of the patients is 8.8 ± 5.8 years. Data on the average age of subjects in each of these subgroups are summarized below (Table 1). 234 patients (55.6 %) were female and 187 (44.4 %) were male (S1 Table).

**Table 1.**
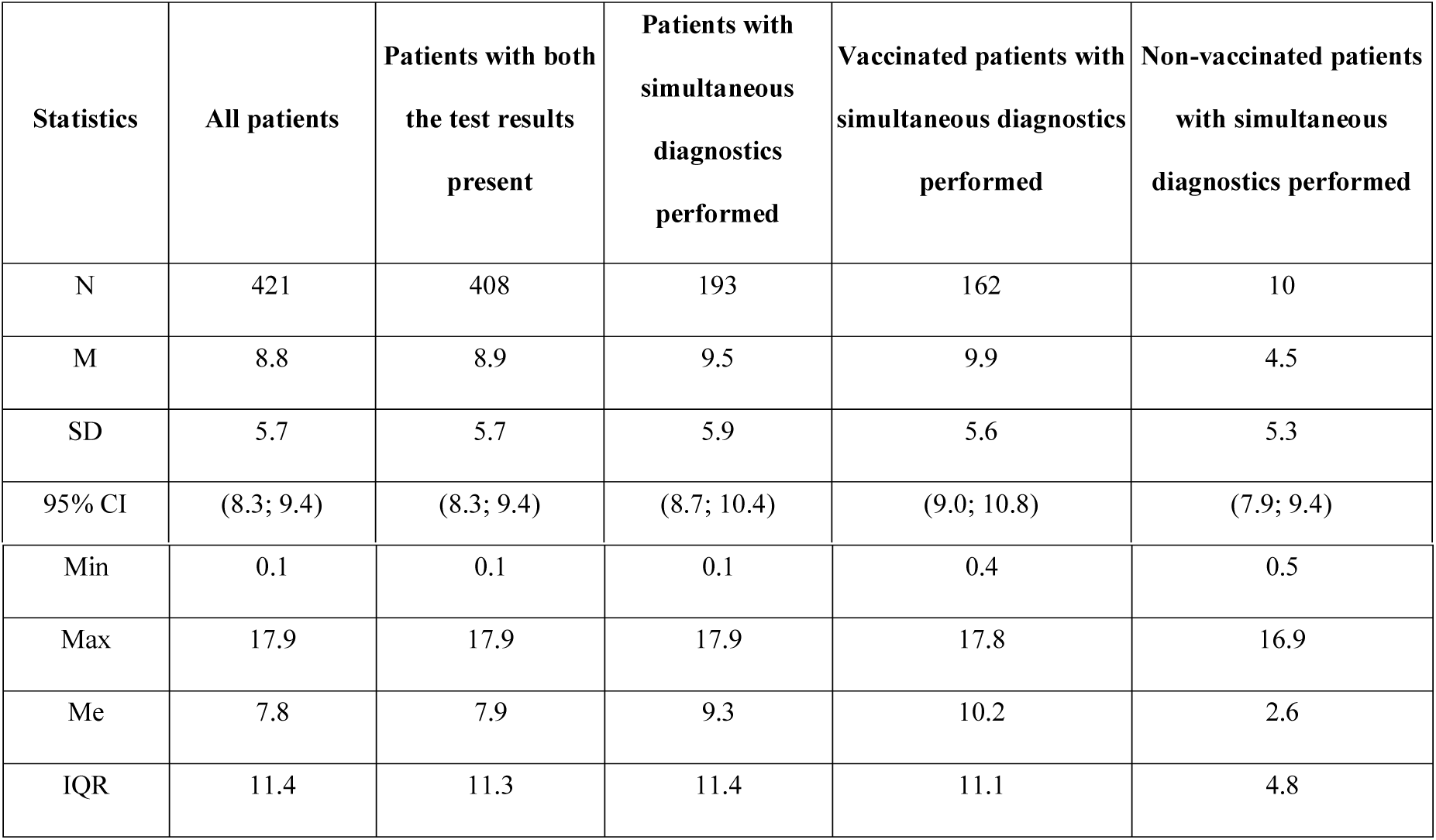
Descriptive statistics of the patients’ age (years), n=421.

| Statistics | All patients | Patients with both the test results present | Patients with simultaneous diagnostics performed | Vaccinated patients with simultaneous diagnostics performed | Non-vaccinated patients with simultaneous diagnostics performed |
| --- | --- | --- | --- | --- | --- |
| N | 421 | 408 | 193 | 162 | 10 |
| M | 8.8 | 8.9 | 9.5 | 9.9 | 4.5 |
| SD | 5.7 | 5.7 | 5.9 | 5.6 | 5.3 |
| 95% CI | (8.3; 9.4) | (8.3; 9.4) | (8.7; 10.4) | (9.0; 10.8) | (7.9; 9.4) |
| Min | 0.1 | 0.1 | 0.1 | 0.4 | 0.5 |
| Max | 17.9 | 17.9 | 17.9 | 17.8 | 16.9 |
| Me | 7.8 | 7.9 | 9.3 | 10.2 | 2.6 |
| IQR | 11.4 | 11.3 | 11.4 | 11.1 | 4.8 |

Out of 421 patients, 408 patients had the results of both tests, 193 patients were simultaneously given 2 tests: TST with 2 TU PPD-L and Diaskintest, 162 of whom were BCG-vaccinated, 10 were not vaccinated, no data were reported for 21, due they were migrants.

The diagnosis of tuberculosis is confirmed by the discovery of the pathogen in a small number of cases (13/421 - 3.1 %). The main diagnosis is based on a combination of clinical and radiological data.

In the overwhelming majority of cases, tuberculosis of the intrathoracic lymph nodes (about 50 %, Table 2) was noted among children and adolescents under 17 years with a confirmed diagnosis of tuberculosis.

**Table 2.** Distribution of the patients of the type of tuberculosis, n=421.

| Type of tuberculosis | All patients | Patients with both the test results present | Patients with simultaneous diagnostics performed | Vaccinated patients with simultaneous diagnostics performed | Non-vaccinated patients with simultaneous diagnostics performed |
| --- | --- | --- | --- | --- | --- |
| Intrathoracic lymph nodes tuberculosis | 200/421 (47.5 %) | 196/408 (48.0 %) | 85/193 (44.0 %) | 71/162 (43.8 %) | 6/10 (60.0 %) |
| Primary tuberculosis complex | 84/421 (20.0 %) | 79/408 (19.4 %) | 33/193 (17.1 %) | 28/162 (17.3 %) | 2/10 (20.0 %) |
| Infiltrative lung tuberculosis | 63/421 (15.0 %) | 62/408 (15.2 %) | 28/193 (14.5 %) | 25/162 (15.4 %) | 1/10 (10.0 %) |
| Focal tuberculosis | 55/421 (13.1 %) | 54/408 (13.2 %) | 36/193 (18.7 %) | 30/162 (18.5 %) | 0/10 (0.0 %) |
| Disseminated tuberculosis | 10/421 (1.9 %) | 9/408 (1.7 %) | 7/193 (2.5 %) | 6/162 (3.1 %) | 1/10 (0.0 %) |
| Tuberculoma | 4/421 (1.0 %) | 4/408 (1.0 %) | 0/193 (0.0 %) | 0/162 (0.0 %) | 0/10 (0.0 %) |
| Cavernous tuberculosis | 2/421 (0.5 %) | 2/408 (0.5 %) | 2/193 (1.0 %) | 0/162 (0.0 %) | 0/10 (0.0 %) |
| Pleurisy of tuberculosis etiology | 3/421 (0.7 %) | 2/408 (0.5 %) | 2/193 (1.0 %) | 2/162 (1.2 %) | 0/10 (0.0 %) |

#### Analyses of the sensitivity of the two tests

Of 421 patients having results of Diaskintest, the negative reaction was found within 8/421 (1.9%) subjects, while 9/421 (2.1%) had induration size of less than 5 mm. Thus, the sensitivity of Diaskintest at cut-off > 0 mm was 98.1% (95% CI 96.2%-99.2%), at cut-off ≥ 5 mm 97.9% (95% CI 96.0%-99.0%, at cut-off ≥ 10 mm – 89.8% (95% CI 86.5%-92.5%, and at cut-off ≥ 15 mm 61m1% (95% CI 56.2%-65.7%) (Table 3).

**Table 3.** Sensitivity of the tests on the sample of all the patients, n = 421.

| Diagnostic test | cut-off > 0 mm | cut-off ≥ 5 mm | cut-off ≥ 10 mm | cut-off ≥ 15 mm |
| --- | --- | --- | --- | --- |
| Diaskintest (n=421) | 413/421 (98.1%) | 412/421 (97.9%) | 378/421 (89.8%) | 257/421 (61.1%) |
| 95% CI | 96.2%-99.2% | 96.0%-99.0% | 86.5%-92.5% | 56.2%-65.7% |
| TST (n=414) | 409/414 (98.8%) | 406/414 (98.1%) | 368/414 (88.9%) | 191/414 (46.1%) |
| 95% CI | 97.2%-99.6% | 96.2%-99.2% | 85.5%-91.7% | 41.3%-51.1% |

TST results were available for 414 patients (the parents of the rest have refused to perform TST having done only the Diaskintest). Absence of induration has been revealed for 5/414 (1.2%) subjects, while 8/414 (1.9%) had induration of less than 5 mm. Thus, the sensitivity of TST was 98.8% (95% CI 97.2%-99.6%), 98.1% (95% CI 96.2%-99.2%), 88.9% (95% CI 85.5%-91.7%), and 46.1% (95% CI 41.3%-51.1%), respectively (Table 3). Sensitivity at cut-off ≥ 15 mm was statistically significant lower compared to that of Diaskintest (<0.0001).

408 of 421 subjects had the results of both the tests (Table 4). It was found that at cut-off ≥5 mm the sensitivity of both the tests was similar: Diaskintest 98.3% (95% CI 97.0-99.6%) and TST 98.0% (95% CI 96.7-99.4%), while at cut-off ≥ 10 mm the sensitivity somewhat decrease to 90.0% (95% CI 87.0-93.0%) for Diaskintest and to 88.7% (95% CI 85.6-91.9%) for TST. However, at cut-off ≥ 15 mm the drop in sensitivity was both statistically and clinically significant (p< 0.0001): to 61.5% (95% CI 56.7-66.3%) for Diaskintest and even worse for TST – to 46.3% (95% CI 41.4-51.3%) (Fig 1).

**Table 4.**
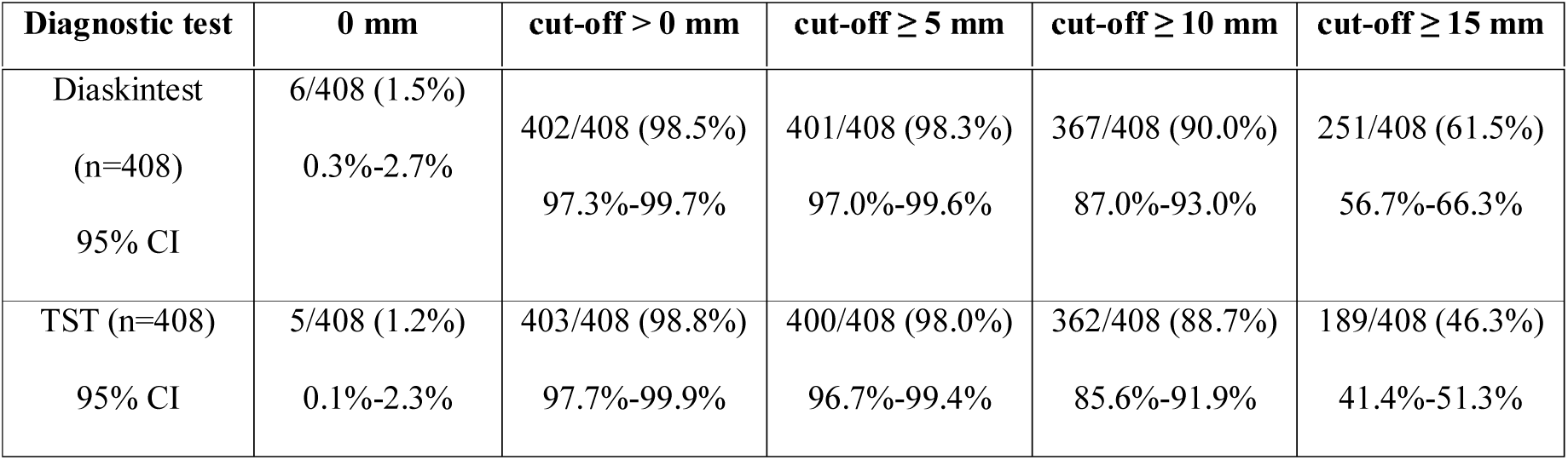
Sensitivity of the tests among the patients having both the tests results, n=408.

**Fig 1.**
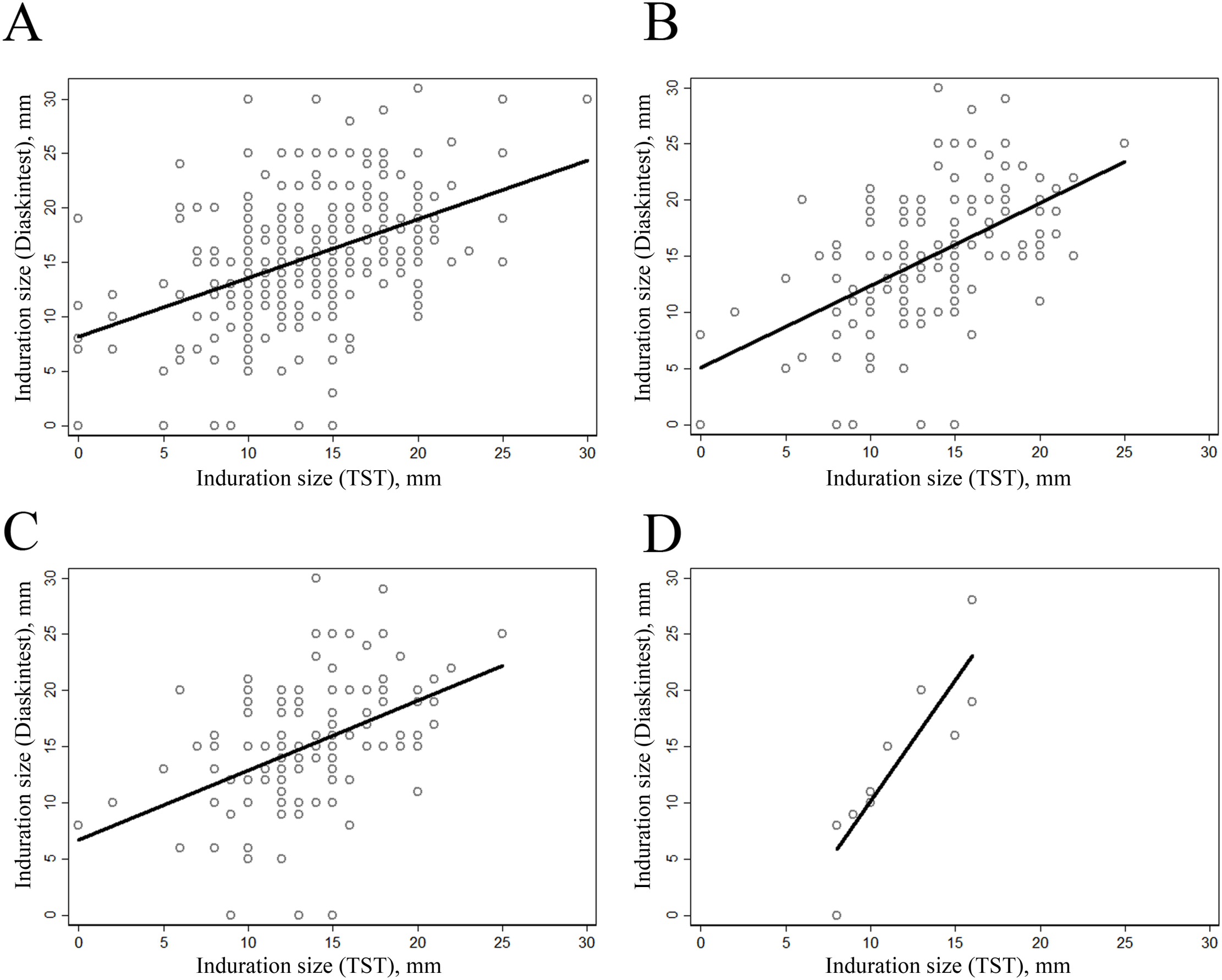
Sensitivity of the two diagnostics tests (a point estimation along with 95% CI): A - the patients with the both results present (n = 408), B - patients with simultaneous diagnostics performed (n = 193), C - vaccinated patients with simultaneous diagnostics performed (n = 162), D - non-vaccinated patients with simultaneous diagnostics performed (n = 10).

The simultaneous diagnostics with both the tests was performed for 193 subjects (Table 5, Fig 1). The results for those patients were very close to the results for the above-mentioned 408 subjects.

**Table 5.** Sensitivity of the tests among the patients with simultaneous diagnostics performed, n=193.

| Diagnostic test | 0 mm | cut-off > 0 mm | cut-off ≥ 5 mm | cut-off ≥ 10 mm | cut-off ≥ 15 mm |
| --- | --- | --- | --- | --- | --- |
| Diaskintest<br>95% CI | 5/193 (2.6%)<br>0.8%-5.9% | 188/193 (97.4%)<br>94.1%-99.2% | 188/193 (97.4%)<br>94.1%-99.2% | 172/193 (89.1%)<br>83.9%-93.2% | 116/193 (60.1%)<br>53.1%-67.2% |
| TST<br>95% CI | 2/193 (1.0%)<br>0.1%-3.7% | 191/193 (99.0%)<br>96.3%-99.9% | 190/193 (98.5%)<br>95.5%-99.7% | 172/193 (89.1%)<br>83.8%-93.1% | 89/193 (46.1%)<br>38.9%-53.4% |

Among patients with the simultaneous using of the two different skin tests, 162 persons (94.2%) were known to be BCG vaccinated (Table 6, Fig 1). Sensitivity of both the tests was similar to that of the sample collected 193 patients, as almost all of them were vaccinated.

**Table 6.** Sensitivity of the tests among the vaccinated patients with simultaneous diagnostics performed, n=162.

| Diagnostic test | cut-off > 0 mm | cut-off ≥ 5 mm | cut-off ≥ 10 mm | cut-off ≥ 15 mm |
| --- | --- | --- | --- | --- |
| Diaskintest | 159/162 (98.2%) | 159/162 (98.2%) | 148/162 (91.4%) | 100/162 (61.7%) |
| 95% CI | 94.7%-99.6% | 94.7%-99.6% | 85.9%-95.2% | 53.8%-69.2% |
| TST | 161/162 (99.4%) | 160/162 (98.8%) | 148/162 (91.4%) | 76/162 (46.9%) |
| 95% CI | 96.6%-100.0% | 95.6%-99.9% | 85.9%-95.2% | 39.0%-54.9% |

As BCG vaccination is mandatory in the newborn and revaccination – in children at the age of 7 years (with negative Mantoux reactions with 2 TU PPD-L), the number of non-vaccinated patients was only 10 subjects, which explains the extremely wide confidence intervals observed.

The comparison of the results between TST and Diaskintest on the sample of patients with the both diagnostics present was performed using McNemar test at different cut-off levels (Table 7). Given the high concordance of the test results, the kappa coefficient, adjusted for the probability of random coincidence of the results was not so high (kappa <0.4), while no statistically significant difference between the two test results was revealed when an equal cutoff level (either ≥ 5 mm or ≥ 10 mm) was applied for both the tests (p > 0.05). It is also interesting to note, that at cut-off ≥ 15 mm Diaskintest was 2.7 times more likely (p < 0.0001) to show a true positive result (OR = 0.37 with 95% CI 0.25-0.55).

**Table 7.**
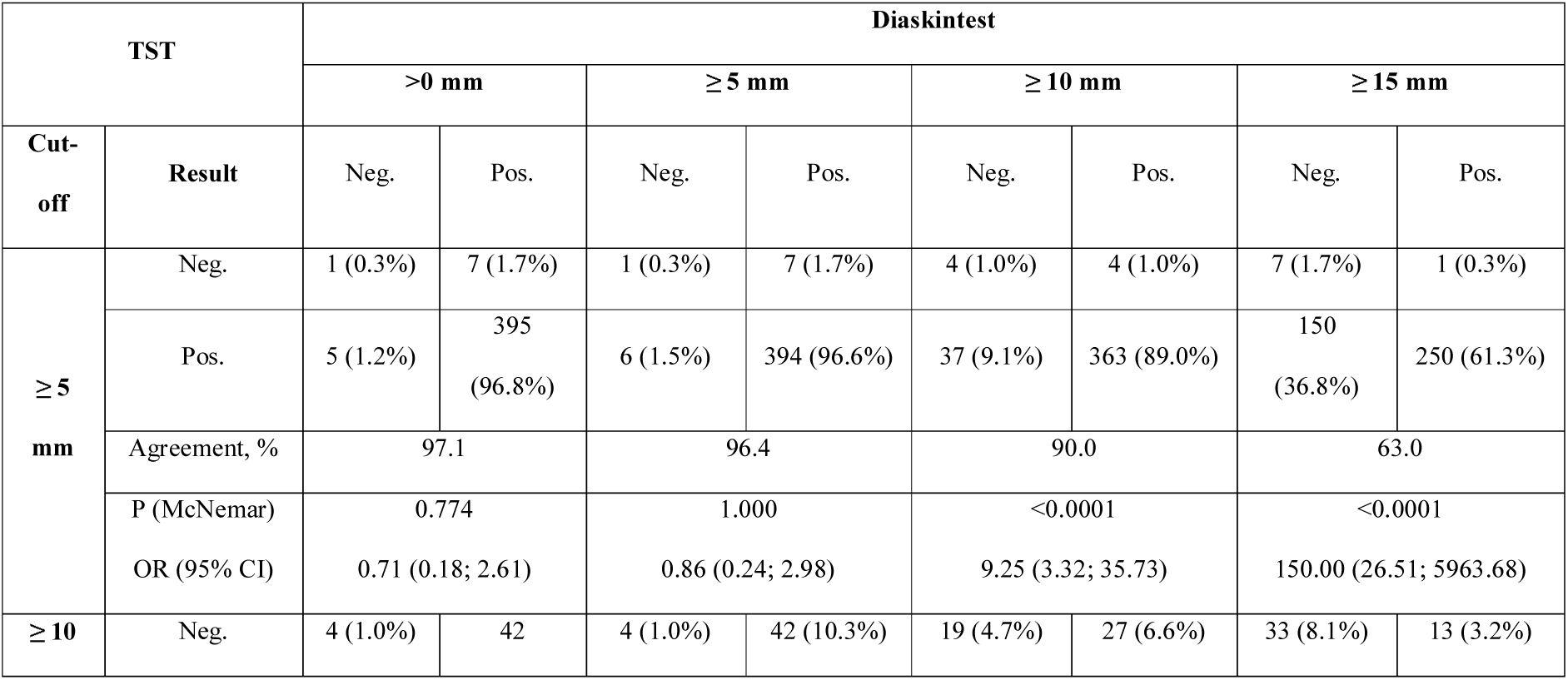
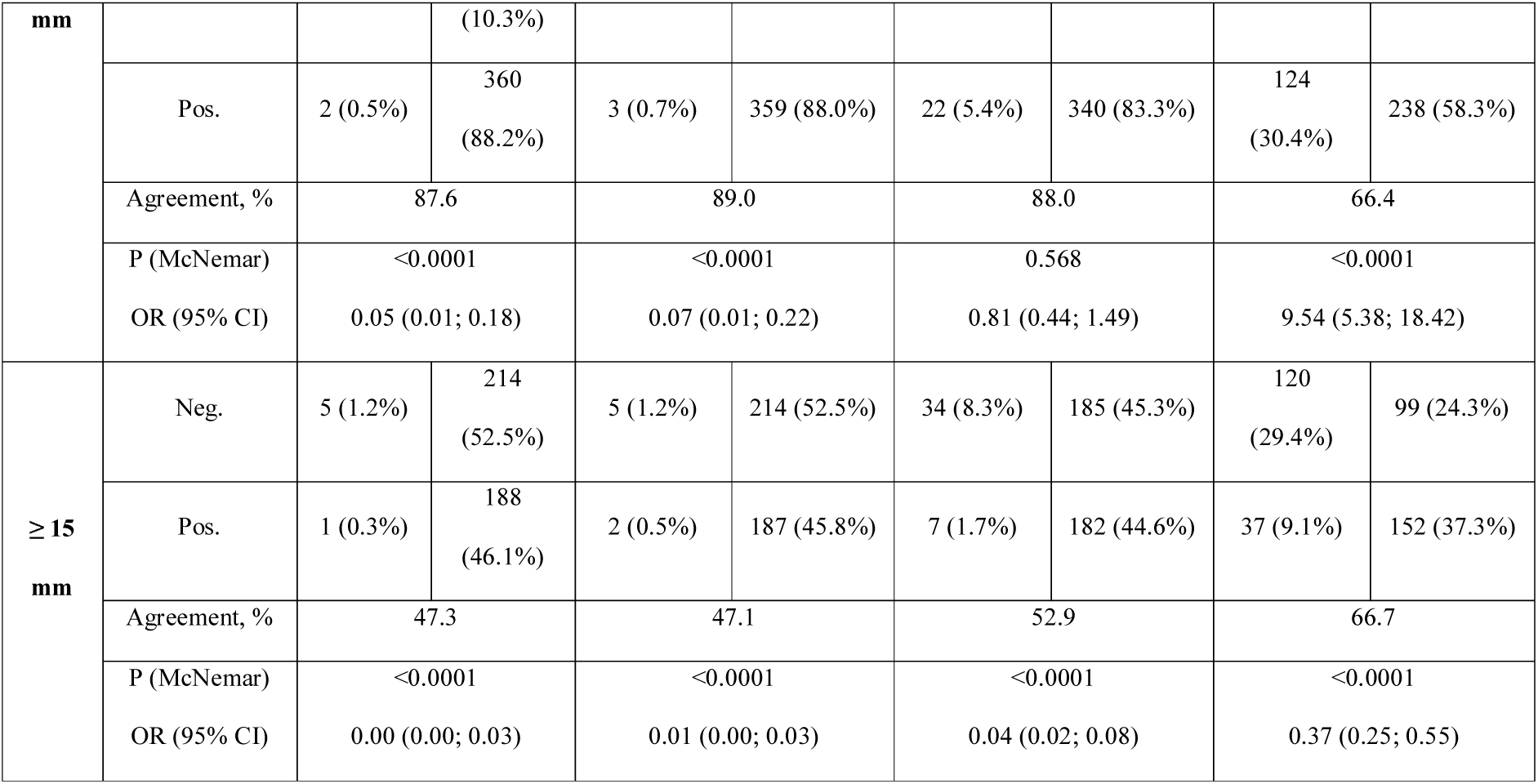
Comparison of the results between TST and Diaskintest at different cut-off levels on the sample of patients with the both results present, n = 408.

Similar results were obtained for the test of patients with simultaneous diagnostics performed (Table 8). No statistically significant difference between the two test results was revealed when an equal cut-off level (either ≥ 5 mm or ≥ 10 mm) was applied for both the tests (p > 0.05). It is also interesting to note, that at cut-off ≥ 15 mm Diaskintest was 2.9 times more likely (p = 0.0004) to show a positive results (OR = 0.34 with 95% CI 0.17-0.64). In subjects with simultaneous tests with TST cut-off ≥ 10 mm and Diaskintest cut-off ≥ 0 mm, the chances of obtaining a positive result of Diaskintest were at least 2.17 times higher (OR = 0.11 with 95% CI 0.01–0.46).

**Table 8.** Comparison of the results between TST and Diaskintest at different cut-off levels on the sample of patients with simultaneous diagnostics performed, n = 193.

| TST |  | Diaskintest |  |  |  |  |  |  |  |
| --- | --- | --- | --- | --- | --- | --- | --- | --- | --- |
| | | >0 mm | | $\geq 5$ mm | | $\geq 10$ mm | | $\geq 15$ mm | |
| Cut-off | Result | Neg. | Pos. | Neg. | Pos. | Neg. | Pos. | Neg. | Pos. |
| ≥ 5 mm | Neg. | 1 (0.5%) | 2 (1.0%) | 1 (0.5%) | 2 (1.0%) | 2 (1.0%) | 1 (0.5%) | 3 (1.6%) | 0 (0.0%) |
|  | Pos. | 4 (2.1%) | 186<br>(96.4%) | 4 (2.1%) | 186<br>(96.4%) | 19 (9.8%) | 171<br>(88.6%) | 74 (38.3%) | 116<br>(60.1%) |
|  | Agreement, % | 96.9 |  | 96.9 |  | 89.6 |  | 61.7 |  |
|  | P (McNemar) | 0.688 |  | 0.688 |  | <0.0001 |  | <0.0001 |  |
|  | OR (95% CI) | 2.00 (0.29; 22.11) |  | 2.00 (0.29; 22.11) |  | 19.00 (3.02; 789.46) |  | - (19.56; -) |  |
| **≥ 10 mm** | Neg. | 3 (1.6%) | 18 (9.3%) | 3 (1.6%) | 18 (9.3%) | 11 (5.7%) | 10 (5.2%) | 17 (8.8%) | 4 (2.1%) |
|  | Pos. | 2 (1.0%) | 170 (88.1%) | 2 (1.0%) | 170 (88.1%) | 10 (5.2%) | 162 (83.9%) | 60 (31.1%) | 112 (58.0%) |
|  | Agreement, % | 89.7 |  | 89.7 |  | 89.6 |  | 66.8 |  |
|  | P (McNemar) | 0.0004 |  | 0.0004 |  | 1.000 |  | <0.0001 |  |
|  | OR (95% CI) | 0.11 (0.01; 0.46) |  | 0.11 (0.01; 0.46) |  | 1.00 (0.37; 2.68) |  | 15.00 (5.56; 56.84) |  |
| **≥ 15 mm** | Neg. | 4 (2.1%) | 100 (51.8%) | 4 (2.1%) | 100 (51.8%) | 19 (9.8%) | 85 (44.0%) | 63 (32.6%) | 41 (21.2%) |
|  | Pos. | 1 (0.5%) | 88 (45.6%) | 1 (0.5%) | 88 (45.6%) | 2 (1.0%) | 87 (45.1%) | 14 (7.3%) | 75 (38.9%) |
|  | Agreement, % | 47.7 |  | 47.7 |  | 54.9 |  | 71.5 |  |
|  | P (McNemar) | <0.0001 |  | <0.0001 |  | <0.0001 |  | 0.0004 |  |
|  | OR (95% CI) | 0.01 (0.00; 0.06) |  | 0.01 (0.00; 0.06) |  | 0.02 (0.00; 0.09) |  | 0.34 (0.17; 0.64) |  |

The results obtained for the BCG vaccinated patients with simultaneous diagnostics performed (**Table 9**) were also similar. While no statistically significant difference between the two test results was revealed when an equal cut-off level (either ≥ 5 mm or ≥ 10 mm) was applied for both the tests (p > 0.05). It is also interesting to note, that at cut-off ≥ 15 mm Diaskintest was 2.8 times more likely (p =0.0009) to show a true positive result (OR = 13/37 = 0.35 with 95% CI 0.17-0.68).

**Table 9.** Comparison of the results between TST and Diaskintest at different cut-off levels on the sample of vaccinated patients with simultaneous diagnostics performed, n = 162.

| TST |  | Diaskintest |  |  |  |  |  |  |  |
| --- | --- | --- | --- | --- | --- | --- | --- | --- | --- |
| | | >0 mm | | $\geq 5$ mm | | $\geq 10$ mm | | $\geq 15$ mm | |
| Cut-off | Result | Neg. | Pos. | Neg. | Pos. | Neg. | Pos. | Neg. | Pos. |
| **≥ 5 mm** | Neg. | 0 (0.0%) | 2 (1.2%) | 0 (0.0%) | 2 (1.2%) | 1 (0.6%) | 1 (0.6%) | 2 (1.2%) | 0 (0.0%) |
|  | Pos. | 3 (1.9%) | 157 (96.9%) | 3 (1.9%) | 157 (96.9%) | 13 (8.0%) | 147 (90.7%) | 60 (37.0%) | 100 (61.7%) |
|  | Agreement, % | 96.9 |  | 96.9 |  | 91.3 |  | 62.9 |  |
|  | P (McNemar) | 1.000 |  | 1.000 |  | 0.0018 |  | <0.0001 |  |
|  | OR (95% CI) | 1.50 (0.17; 17.96) |  | 1.50 (0.17; 17.96) |  | 13.00 (1.95; 552.47) |  | - (15.77; -) |  |
| **≥ 10 mm** | Neg. | 1 (0.6%) | 13 (8.0%) | 1 (0.6%) | 13 (8.0%) | 5 (3.1%) | 9 (5.6%) | 10 (6.2%) | 4 (2.5%) |
|  | Pos. | 2 (1.2%) | 146 (90.1%) | 2 (1.2%) | 146 (90.1%) | 9 (5.6%) | 139 (85.8%) | 52 (32.1%) | 96 (59.3%) |
|  | Agreement, % | 90.7 |  | 90.7 |  | 88.9 |  | 65.5 |  |
|  | P (McNemar) | 0.0074 |  | 0.0074 |  | 1.000 |  | <0.0001 |  |
|  | OR (95% CI) | 0.15 (0.02; 0.68) |  | 0.15 (0.02; 0.68) |  | 1.00 (0.35; 2.84) |  | 13.00 (4.78; 49.50) |  |
| **≥ 15 mm** | Neg. | 2 (1.2%) | 84 (51.9%) | 2 (1.2%) | 84 (51.9%) | 12 (7.4%) | 74 (45.7%) | 49 (30.3%) | 37 (22.8%) |
|  | Pos. | 1 (0.6%) | 75 (46.3%) | 1 (0.6%) | 75 (46.3%) | 2 (1.2%) | 74 (45.7%) | 13 (8.0%) | 63 (38.9%) |
|  | Agreement, % | 47.5 |  | 47.5 |  | 53.1 |  | 69.2 |  |
|  | P (McNemar) | <0.0001 |  | <0.0001 |  | <0.0001 |  | 0.0009 |  |
|  | OR (95% CI) | 0.01 (0.00; 0.07) |  | 0.01 (0.00; 0.07) |  | 0.03 (0.00; 0.10) |  | 0.35 (0.17; 0.68) |  |

The results for non-vaccinated patients with simultaneous tests were not statistically significant due to a small sample size (n = 10) (S2 Table).

#### The size of indurations

Most of the patients had the size of induration ≥10 mm for both the diagnostic tests. At the same time, the percent of patients having induration ≥15 mm on Diaskintest test was higher compared to TST (Fig 2): 61.5% vs 46.3% (for all the patients with the both results present, n=408), 60.1% vs 46.1% (for the patients with simultaneous diagnostics performed, n=193), 61.7% vs 46.9% (for the vaccinated patients with simultaneous diagnostics performed, n = 162), and 30.0% vs 50.0% (for the non-vaccinated patients with simultaneous diagnostics performed, n = 10).

**Fig 2.**
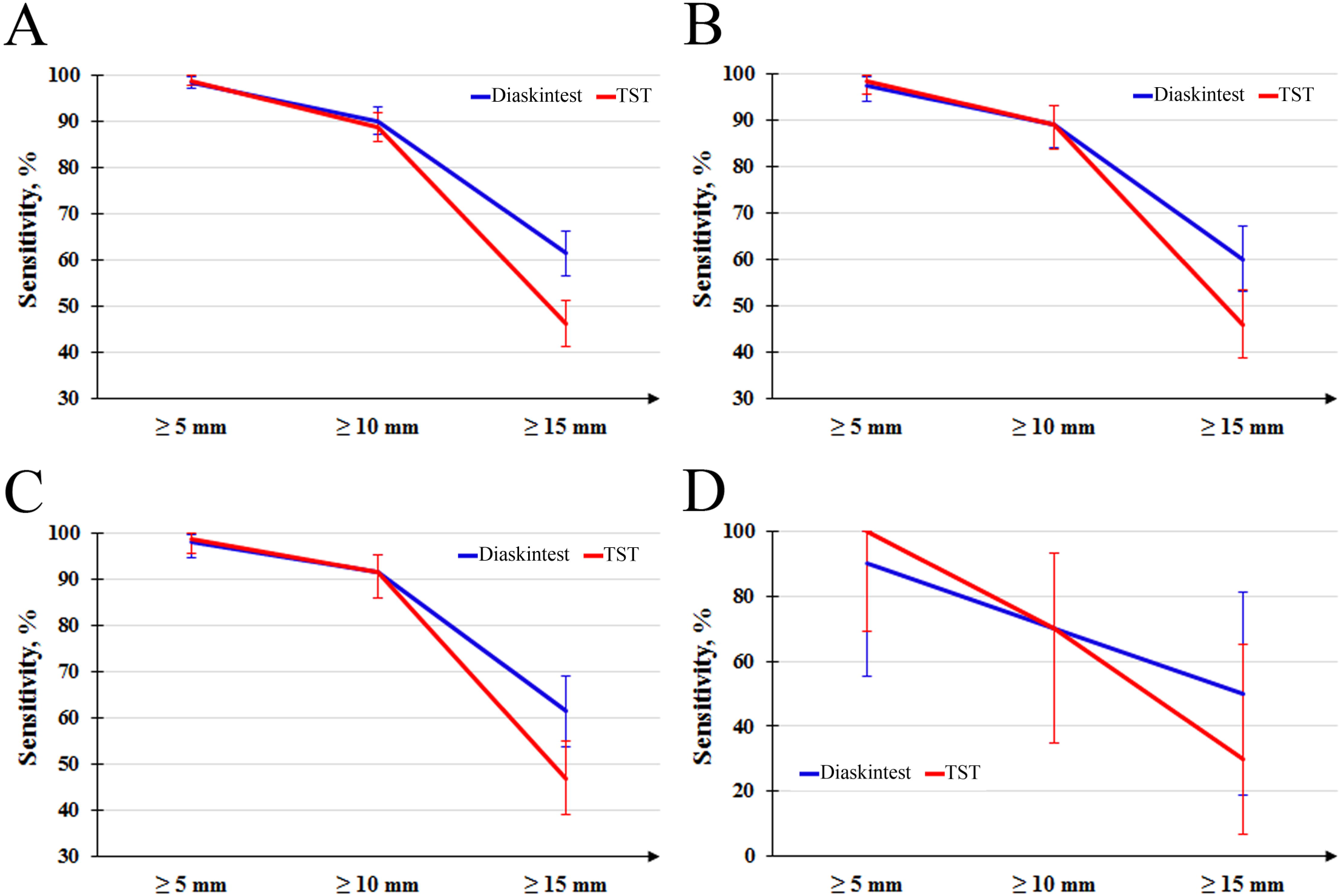
Induration size for the two diagnostics tests: A - the patients with the both results present (n = 408), B - patients with simultaneous diagnostics performed (n = 193), C - vaccinated patients with simultaneous diagnostics performed (n = 162), D - non-vaccinated patients with simultaneous diagnostics performed (n = 10).

Generally, the patients in all the samples have shown a higher induration size for Diaskintest compared with TST. The analysis performed with a paired t-test has shown statistically significant differences (p < 0.001) except for the non-vaccinated patients with simultaneous diagnostics performed, where only a small sample size was available (Fig 3, S1 Fig 3). The difference was on average 1.9 mm (with 95% CI 1.4-2.4 mm) for all the patients with both the test results present. Similar effect size was found for the patients with simultaneous diagnostics performed: on average 1.4 mm (with 95% CI 0.8-2.1 mm). Despite for the non-vaccinated patients the difference has not reached a level of statistical significance, the point estimate was also 2.0 mm (13.6 ± 7.8 mm for Diaskintest versus 11.6 ± 3.2 mm for TST).

**Fig 3.**
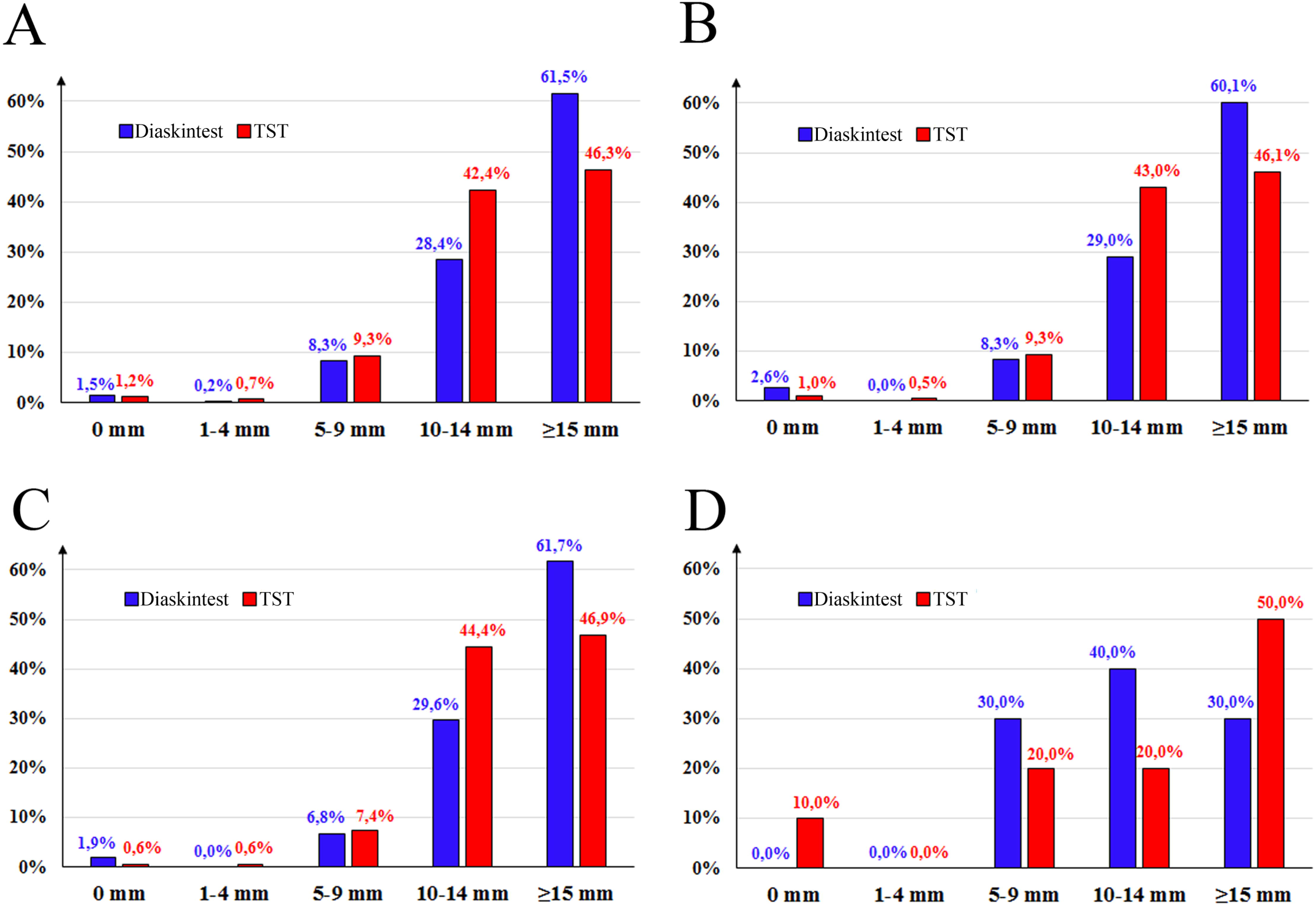
Boxplot for the induration sizes of the two diagnostics tests: A - the patients with the both results present (n = 408), B - patients with simultaneous diagnostics performed (n = 193), C - vaccinated patients with simultaneous diagnostics performed (n = 162), D - non-vaccinated patients with simultaneous diagnostics performed (n = 10).

The Pearson correlation between induration sizes of the two tests was of clinical and statistical significance (above 0.4, p < 0.001) in all the samples with maximum of 0.878 observed for the non-vaccinated patients with simultaneous diagnostics performed (Fig 4).

**Fig 4.**
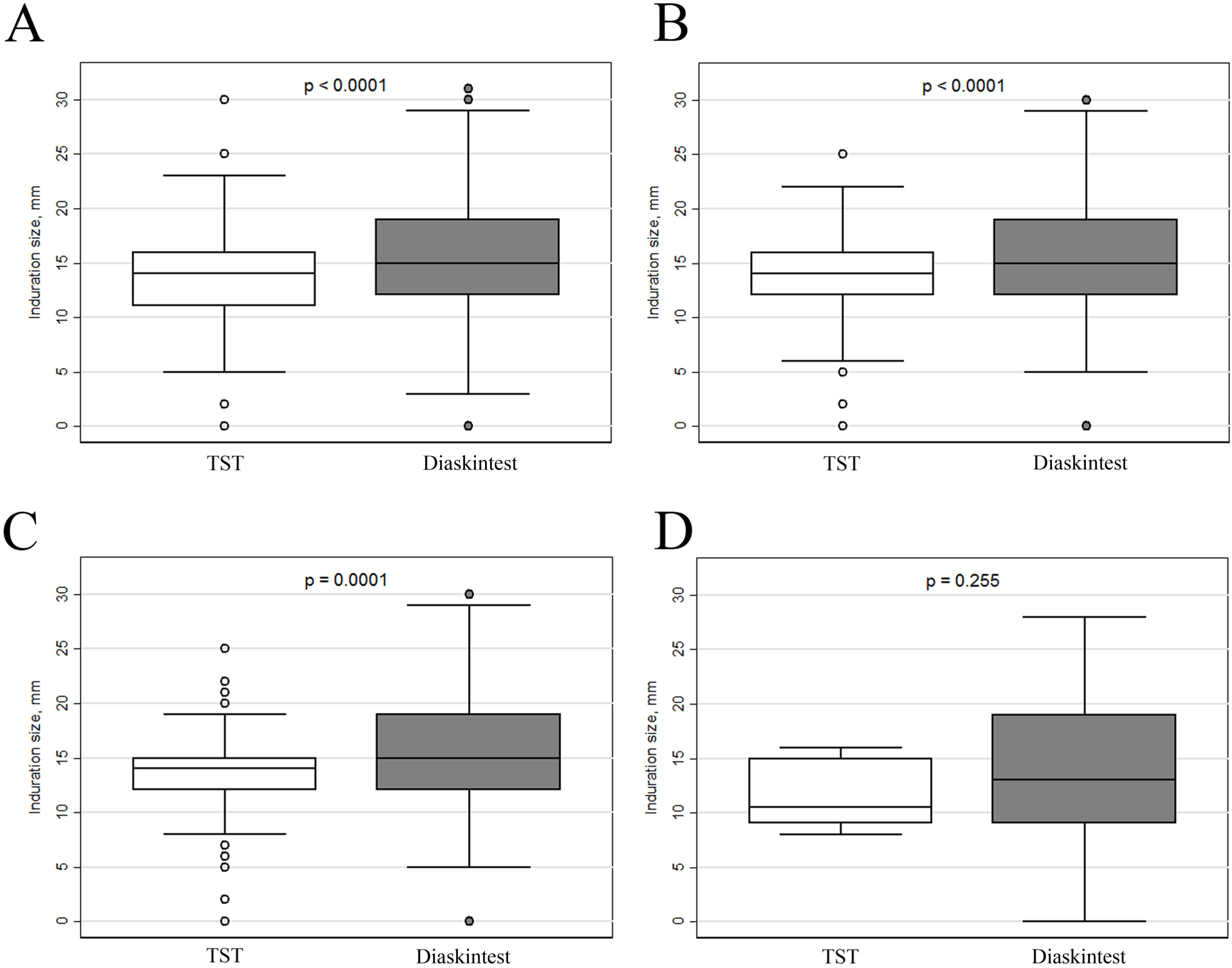
Scatter-plot for the induration sizes of the two diagnostics tests: A - the patients with the both results present (n = 408), B - patients with simultaneous diagnostics performed (n = 193), C - vaccinated patients with simultaneous diagnostics performed (n = 162), D - non-vaccinated patients with simultaneous diagnostics performed (n = 10).

The comparative study of the sensitivity of TST and Diaskintest in the same of 408 children with newly diagnosed tuberculosis of respiratory organs in different cut-offs demonstrated the highest sensitivity of TST and Diaskintest for cut-off ≥ 5 mm: Diaskintest 98.3 % (95 % CI 97.0–99.6 %), TST 98.0 % (95 % CI 96.7–99.4 %), with cut-off ≥10 mm, the sensitivity of tests decreases to Diaskintest insignificantly to TST to a greater extent: 90.0 % (95 % CI 87.0–93.0 %), and 88.7 % (95 % CI 85.6–91.9 %), respectively; but with cut-off ≥ 15 mm, the decrease in sensitivity is statistically significant: 61.5 % (95 % CI 56.7–66.3 %) to Diaskintest, and to TST 46.3 % (95 % CI 41.4–51.3 %), p <0.0001.

Discordant results with negative Diaskintest were observed: in 2013, in 2 children aged 2 and 3 months who were born by women with tuberculosis. Children were examined as contacts – both tests were negative. Another child suffered from chronic adrenal insufficiency (TST (+), Diaskintest (-)), he was examined as contact with a tuberculous father and had disseminated pulmonary tuberculosis; another child with disseminated pulmonary tuberculosis and one with focal pulmonary tuberculosis (Mantoux test (+), Diaskintest (-)). In 2014, one teenager with pleurisy (Mantoux test (+), Diaskintest (-)), in 2015, 4 children with HIV infection and disseminated pulmonary tuberculosis (Mantoux test (-), Diaskintest (-)); 2 teenagers with infiltrative pulmonary tuberculosis (Mantoux test (+), Diaskintest (-)), 1 child with hilar tuberculosis in the calcification phase (Mantoux test (+), Diaskintest (-)), in 2016, 1 child, 6 months old, not vaccinated with BCG, with disseminated tuberculosis had a negative reaction (Mantoux test (+), Diaskintest (-)).

## Discussion

In Russia, BCG vaccination of newborns and revaccination at 7 years (with negative TST with 2 TU PPD-L) is mandatory. Mantoux tuberculin test with 2 TU PPD-L was also mandatory so far. It is clear that under such conditions the specificity of the sample is extremely low. The situation is aggravated by the fact that in Russia there is one limit of positive results – 5 mm for all – vaccinated and unvaccinated. At the same time, positive reactions to the Mantoux test in children and adolescents are registered in 75 %, and the increase in the reaction is observed annually only in 0.9 % of the tuberculin-positive ones. Annually, the rate of primarily infected children among the tuberculin-positive is 0.67 %, and 0.06 % in adolescents [41]. In such conditions, there was a need for a specific test capable of distinguishing post-vaccination allergy. Such a test was a skin test with a tubercular recombinant allergen (Diaskintest) containing ESAT-6–CFP-10 protein. [42]. Since 2009, at the order of the Ministry of Health of Russia, the Diaskintest has been used for differential diagnosis of postvaccinal and infectious allergies. All children with a positive reaction to Diaskintest should be examined to identify local forms of tuberculosis and, in the absence of such, to receive preventive treatment for LTBI. Detection of tuberculosis among people with positive reactions to Diaskintest is ten times higher than among tuberculin-positive [43].

The comparative study of the sensitivity of Mantoux test and Diaskintest in the same of 408 children with newly diagnosed respiratory tuberculosis at different cut-offs demonstrated the highest sensitivity of Mantoux and Diaskintest skin tests for cut-off ≥ 5 mm (98.3 %), but with cut-off ≥ 15 mm, the decrease in sensitivity is statistically significant Diaskintest, and to TST, p <0.0001 Thus, the optimal cut-off of 5 mm for the test with Diaskintest is shown. With such a cut-off, the specificity of the Mantoux test is extremely low, and the Diaskintest is optimal.

In phase III clinical studies of C-tb Danish skin test, three tests were compared: QuantiFERON-TB Gold In-Tube^®^, tuberculin test (TST), and C-tb skin test. The results showed that C-Tb and QFT were consistent in 785 of 834 (94 %). Sensitivity of C-Tb was 69 % in *TB*, which is 14 % lower than that of QFT. The comparison of C-Tb, TST, and QFT showed that the TST specificity in BCG vaccinated patients increased from 62 % to 92 %, and the consistency of TST with QFT and C-Tb increased to 90 % only at the ≥15 mm cut-off of TST positive result in BCG vaccinated patients; the specificity of QFT and C-Tb is 96 % [*38*].

We compared the sensitivity results obtained with Diaskintest and the results of the skin test with C-tb [*38*] and let us assume the reasons for the lower sensitivity of the latter: firstly, adults with lower sensitivity compared to children were examined, and secondly, those who completed the treatment a long time ago were enrolled to the group of those surveyed, and they probably had a negative reaction, as was observed in our studies with Diaskintest [44]; the possibility of insufficient sensitivity in the difference in dosage in C-tb 0.1 μg (Diaskintest – 0.2 μg) is not excluded.

The high sensitivity of Diaskintest in children can be explained by the optimal diagnosis of the disease, which is facilitated by the fact that children from 1 year of age are subject to annual tuberculin diagnostics, the moment of primary infection is fixed and they are subject to examination at the phthisiatrician. Our comparative studies of the sensitivity of Diaskintest and QuantiFERON-TB Gold In-Tube^®^ tests showed high sensitivity of not only Diaskintest, but also QuantiFERON-TB Gold In-Tube^®^ (more than 90 %) [45], which can be explained by the adequate diagnosis of tuberculosis in children, which is a difficult task in the absence of clinical manifestations and bacterial excretion. After the introduction of Diaskintest to wide practice in 2010 and the prescription of preventive therapy to children with a positive reaction, the incidence of tuberculosis in Moscow in 2016 decreased from 10.6 to 5.5. The incidence in adolescents showed a decrease from 28.7 to 11.6, with the incidence of the resident population almost 2 times lower in both children and adolescents [46]. An important role is played by the fact that the detection, diagnosis and treatment of tuberculosis and LTBI in children is fully funded, regardless of the social belonging of children, their citizenship and their permanent place of residence. All undergo the vaccination, annual tuberculin diagnostics, Diaskintest (with positive reactions to TST), computed tomography with positive Diaskintest reaction, bacteriological and molecular genetic examination, long-term controlled therapy in a hospital with tuberculosis, and outpatient or in a sanatorium with LTBI (at the request of parents). Data on the high sensitivity of the skin test with the drug containing the specific ESAT6–CFP10 protein, corresponds to the data of S. Sollai [29] that in high-income countries the sensitivity of IGRA tests is higher than in low-income countries.

Diaskintest test is simple to perform, it is currently used in primary health care, it is cheap (the cost is similar to the Mantoux test) and is now implemented by the order of the Ministry of Health into the practice of Russia as a screening method instead of Mantoux test in children over 7 years old.

## Conclusion

In children and adolescents with tuberculosis, Diaskintest at a dose of 0.2 µg/ml and the Mantoux test with 2 TU PPD-L have a high sensitivity (98%) with a cut-off of 5 mm; however, at cut-off ≥ 15 mm sensitivity is significantly reduced, and the decrease is more pronounced in the Mantoux test. The advantage of Diaskintest is that, unlike the Mantoux test, it has high specificity under the conditions of mass BCG vaccination. The test is cost-effective, simple to carry out, and can be used in mass screening.

## Conflict of interest

The authors declare no conflict of interest.

## Acknowledgement

The study had no sponsorship.

## Supporting information

**S1 Table. Gender distribution of the patients, n= 421.**

**S2 Table. Comparison of the results between TST and Diaskintest at different cut-off levels on the sample of non-vaccinated patients with simultaneous diagnostics performed (n = 10).**

**S1 Fig 3. Descriptive statistics of the induration size (mm) of the Diaskintest (n=421) and TST, n=414.**

